# The Periaqueductal Gray Selectively Supports Reversal Learning During a Flexible Discrimination Task in Mice

**DOI:** 10.64898/2026.01.19.700312

**Authors:** Daniela Lichtman, Eyal Bergmann, Jonathan Nicholas, Raphael T. Gerraty, Itamar Kahn

**Affiliations:** Department of Neuroscience, Rappaport Faculty of Medicine, Technion – Israel Institute of Technology, Haifa 31096, Israel; Department of Neuroscience, Mortimer B. Zuckerman Mind Brain Behavior Institute, Columbia University, New York, NY; Department of Psychology, New York University, New York, NY

## Abstract

Flexible, goal-directed behavior depends on the ability to update value representations in response to changing contingencies. This ability depends on distributed brain networks, calling for use of whole-brain imaging. Widely used in human research, whole-brain imaging in rodents has been a major challenge that has only recently been properly addressed. Using functional MRI in behaving mice performing a go/no-go odor discrimination task, we compared neural activity during initial cue-reward learning (acquisition) and subsequent contingency reversal. To link neural activity to underlying learning processes, we modeled value updating using a model-free reinforcement-learning algorithm. Trial-by-trial estimates of state-action values allowed us to dissociate acquisition from reversal-related signals, revealing that ventral striatal responses tracked expected value during acquisition, whereas reversal learning additionally recruited the periaqueductal gray (PAG), a midbrain structure classically linked to threat processing and aversive learning. PAG activity closely followed model-derived signatures of reversal learning, implicating it in the suppression of previously rewarded actions and in updating behavior in the absence of explicit punishment. These findings reveal a previously unrecognized computational role for the PAG in value-based decision-making and cognitive flexibility, and substantiate task-fMRI as a powerful tool to study the rodent brain at a mesoscale resolution.

## Introduction

Decision-making often occurs under conditions of uncertainty, yet individuals are generally capable of making decisions that lead to beneficial outcomes. This capacity relies on neural mechanisms that support the estimation of outcomes and the adjustment of behavior in response to changing environmental conditions, a process referred to as cognitive flexibility (Diamond, 2013). Reversal learning paradigms are prominently used to study cognitive flexibility across species (Highgate & Schenk, 2020; Izquierdo et al., 2017), in which switching of contingencies creates a violation of learned expectancies, resulting in rapid changes of behavioral responses. This requires the ability to detect unexpected outcomes, suppress previously reinforced responses, and update value representations in the light of new contingencies. These adaptive processes are supported by reinforcement learning mechanisms in the brain.

Reinforcement learning refers to the process by which organisms learn to associate specific stimuli or actions with rewarding or punishing outcomes, adapting their behavior to maximize positive results and minimize negative ones (Dayan & Niv, 2008; O’Doherty et al., 2017). A fundamental pathway supporting reinforcement learning is the dopaminergic input from the ventral tegmental area (VTA) to the ventral striatum, and specifically the nucleus accumbens, which serves as a teaching signal and plays a critical role in motivation and value-based decision-making (Barto, 1995; Glimcher, 2011; Montague et al., 1996; O’Doherty, 2004; Salamone & Correa, 2012; Schultz et al., 1997). Yet this mesolimbic circuitry does not operate in isolation. Converging evidence shows that other brain regions, including the prefrontal cortex, hippocampus, amygdala, and potentially even the periaqueductal gray (PAG), shape how reinforcement-learning related signals are generated, interpreted, and used (Ballard et al., 2019; Lee et al., 2012; Muller et al., 2024; Paton et al., 2006; Roy et al., 2014). Fully characterizing the circuitry involved in these behaviors therefore requires whole-brain approaches that can capture how canonical reward pathways interact with systems traditionally studied in other contexts (e.g., in aversion and defense).

Considerable understanding of brain-wide circuits contributing to reinforcement learning processes has come from human fMRI studies, where computational modeling of trial-by-trial neural signals has proven invaluable in dissecting the mechanisms by which brain regions encode learning, predict behavior, and interact within functional networks (Daw et al., 2006; Niv, 2009; O’Doherty et al., 2003, 2007). Yet, despite their insight, human studies are inherently limited in their ability to establish causality, perform precise circuit manipulations, or achieve the level of experimental control possible in rodents. Extending this whole-brain imaging approach to behaving rodents therefore offers a unique opportunity to link systems-level network dynamics with cellular and molecular mechanisms of learning, bridging a critical translational gap. Until recently, most rodent imaging studies focused on resting-state fMRI to map functional connectivity across the mouse brain, further linking it with behavior (Bergmann et al., 2020; Grandjean et al., 2017; Lichtman et al., 2021; Liska et al., 2018). Recent advances by us and others, including the development of awake imaging setups and rodent-specific hemodynamic models (Bergmann et al., 2025; Desai et al., 2011; Fonseca et al., 2020; Han et al., 2019; Kahn et al., 2011; Lawen et al., 2025; Winkelmeier et al., 2022), now enable the implementation of reliable task-based fMRI. These developments open the door to probing not only canonical reward regions but also underappreciated contributors, offering a systems-level perspective that is critical for understanding the distributed computations underlying reinforcement learning and behavioral flexibility, which can then be further investigated and manipulated using invasive tools available in rodents.

Here, we used task-based fMRI of behaving mice to characterize the neural processes involved in a value-based decision-making task. We developed a non-invasive MR-compatible platform enabling high-resolution behavioral monitoring of head-fixed mice, which facilitated a longitudinal study of mice engaged in a go/no-go odor discrimination task followed by rule reversal, allowing us to investigate the distinct neural mechanisms engaged in initial acquisition versus reversal learning. We analyzed fMRI data using subject-specific trial-by-trial parameters derived from a reinforcement-learning model to better capture the temporal dynamics of learning, thereby improving predictive power and accounting for the small sample sizes feasible in rodent studies. This approach revealed the involvement of different brain regions in acquisition and reversal learning, including areas well-established as involved in goal-directed behavior like the nucleus accumbens, the dorsomedial striatum and the orbitofrontal cortex, but also, surprisingly, the PAG, which we found to be involved specifically in the reversal phase. Further examination showed that the PAG exhibits differential activity depending on correct behavioral outcome, is active during inhibition of lick responses to a “no-go” odor and is inactive during lick responses to a “go” odor.

## Materials and Methods

### Ethics

All animal experiments were conducted in accordance with the United States Public Health Service’s Policy on Humane Care and Use of Laboratory Animals and approved by the Institutional Animal Care and Use Committee of the Technion—Israel Institute of Technology.

### Animals and housing conditions

Twelve C57Bl/6J male mice (2–3 months old) were housed in groups of 2–5 animals per cage in a reversed 12-h light–dark cycle with food and water available *ad libitum* prior to water restriction. The housing room was maintained at 23 ± 2 °C. All experiments were conducted during the dark phase.

### Head-post surgery

To minimize head movement during scanning, mice were implanted with MRI-compatible head posts, as previously described (Bergmann et al., 2025; Lichtman et al., 2021). Briefly, mice were anesthetized with isoflurane (1.5–2.5%), the scalp and periosteum were removed from above the surface of the skull, and a head post was attached to the skull using dental cement (C&B Metabond, Parkell, Brentwood, NY, United States). Mice received a subcutaneous injection containing broad-spectrum antibiotics (Cefalexin) and analgesia (Buprenorphine) during the surgery and daily for at least 3 days after the surgery, and were maintained in their home cage for a postoperative recovery period of 1 week.

### MR-compatible behavioral setup

Experiments were conducted using an MRI-compatible behavioral platform designed to enable precise odor delivery, water reward control, and simultaneous recording of sniffing and licking behavior. The system included a custom head-fixation cradle, an air-dilution olfactometer for rapid odor presentation, the design of which has been previously described in detail (Arneodo et al., 2018; Shusterman et al., 2011), a non-invasive sniff sensor, a pressure-based lick detector, and a calibrated water-delivery mechanism. To adapt traditional olfactory setups for the MRI environment, the olfactometer was positioned outside the scanner bore with its output routed through a final solenoid valve located near the head-fixation apparatus. This configuration minimized magnetic interference while maintaining fast odor onset kinetics (steady-state concentration reached within ∼100 ms). Sniffing was measured through a modified odor port connected to a miniature pressure transducer via short capillary tubing, providing stable, artifact-free respiration signals. Lick detection was achieved using a pressure-based MRI-compatible transducer, and water rewards (∼2.5 µl per drop) were delivered through a solenoid-gated line controlled by an Arduino microcontroller. The olfactometer delivered a continuous flow of clean air (992 ml/min) that switched seamlessly to an odorized stream during stimulus presentation, preventing mechanical cues. Odorized air was generated by diverting nitrogen through odorant vials approximately 1 s before valve opening and mixing it into the main airflow to achieve a tenfold dilution. The final odor pulse lasted 1 s, after which clean air resumed. All flow paths were constructed from Teflon to prevent odor contamination. Odorant concentration profiles were verified using a photoionization detector (PID, Aurora Scientific, model 200B). All behavioral and control signals including sniffing, licks, valve triggers, and reward timing were synchronized and recorded via MATLAB-based scripts. This high-resolution monitoring approach provided precise temporal alignment between behavioral events and fMRI acquisition, ensuring accurate characterization of sensory, motor, and reward-related processes. For a more detailed description of the experimental setup, see Bergmann et al. (2025).

### Flexible discrimination learning

Mice underwent an instrumental go/no-go odor discrimination task followed by a rule reversal in which odor contingencies were switched.

Following postoperative recovery, mice were single-housed and placed on a 10-day water restriction schedule, during which the amount of water was gradually reduced from 5 ml to 1 ml (1 ml per day). Water restriction was maintained throughout the experiment, including weekends. During pretraining (3–7 sessions), mice were first habituated to the setup to ensure acclimation to the scanner environment and were trained during mock scanning to lick a spout to obtain water rewards, with increasing inter-reward intervals (3–7 s), to shape stable licking behavior. Once mice consistently licked to consume water in the MRI setup and exhibited stable sniffing signals, they were scanned while performing the go/no-go task. All animals learned the task (i.e., there was no attrition).

The task consisted of multiple six-minute blocks, with 50 trials (25 per odor) per block, in which two neutral odorants (Pinene and Ethyl-Acetate) were presented pseudo-randomly. Each odor signaled either a rewarded (“go”) or unrewarded (“no-go”) condition. On each trial (2.5 s), an odor was presented for 1 s, and the animal was given a 2 s response window from odor onset. Inter-trial interval ranged from 5–12.5 s. Given a correct lick response to the go odor, the reward was delivered immediately. Lick responses to the no-go odor were not explicitly punished (only implicitly as water was not provided for these incorrect responses), and no lick responses were not rewarded.

During *Acquisition*, mice (*n* = 12) underwent 8–16 sessions (**Supplementary Table 1**) in which they learned to produce a lick response to the go odor and to withhold licking to the no-go odor. The first *Acquisition* session included 10–15 min of lick training during scanner calibration, allowing acquisition of fMRI data of the first odor presentation. In subsequent sessions, 1–2 blocks of the task were performed before fMRI data acquisition to allow for scanner calibration. These blocks were included in the behavioral data presented in **Figure 1** but were not included in the fMRI analyses in subsequent figures.

**Figure 1.**
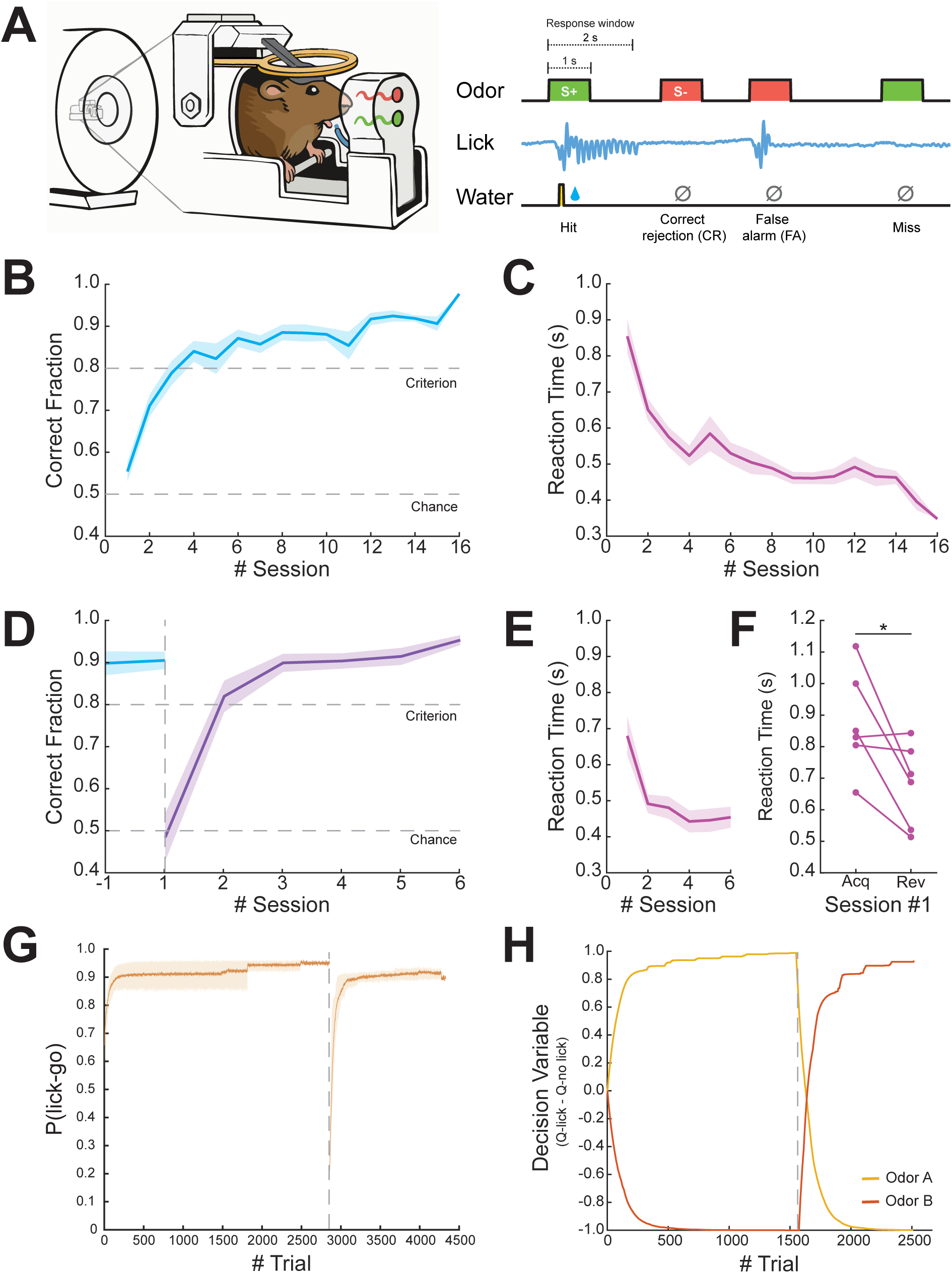
Mice display flexible discrimination learning in a Go/No-go task in an experimental setup allowing for task-based fMRI. **(A)** Illustration of behavioral setup and task design. Odors were presented pseudo-randomly for 1 second with a response window lasting 2 seconds from odor onset. **(B)** Learning curve showing the averaged ratio of correct responses (Hit and Correct Rejection events) during task acquisition (*n* = 12). **(C)** Averaged reaction time of lick responses to Hit trials as a function of session. **(D)** Learning curve showing averaged ratio of correct responses (*n* = 6). Blue line indicates the last acquisition session prior to reversal. Purple line indicates the reversal phase. **(E)** Averaged reaction time of lick responses to Hit trials as a function of sessions during the reversal phase. **(F)** Pairwise comparison of the averaged reaction time for the first session in acquisition vs. reversal. **(G)** Averaged probability of choosing to lick in response to ‘go’ trials (*n* = 6). The gray dashed line indicates the rule reversal onset (95% *CI*_acquisition_ = [0.580, 0.966], 95% *CI*_reversal_ = [0.157, 0.942]). (**H)** Decision variable (Q_lick_ – Q_no-lick_) parameter of a representative animal as computed by the Q-learning model throughout the experiment. The yellow line represents odor A and the orange line represents odor B. The gray dashed line indicates the time of rule reversal. Data are shown as mean ± SEM (B, C, D and E). \**p* < 0.05.

Next, a subset of this cohort (*n* = 6) underwent a *Reversal* phase for 5–6 sessions (**Supplementary Table 1**), during which they learned to lick in response to the presentation of the previously acquired no–go odor and to suppress the previously acquired lick response to the go odor. The *Reversal* group was randomly selected and showed learning performance similar to that of the remaining *Acquisition*-only mice (see results section). In the first *Reversal* session, mice performed 1–2 blocks of the original odor discrimination test during scanner calibration to ensure fMRI data acquisition for the first *Reversal* block. Subsequent sessions were performed using only the *Reversal* task, for which the first 1–2 blocks started before the fMRI data were acquired and therefore contributed to the behavioral data presented in **Figure 1**, but not to fMRI analyses in subsequent figures. Odor identity (go vs. no-go in the *Acquisition* phase) was counterbalanced across animals.

### Reinforcement learning model

In order to model reinforcement learning we used a previously described (Nicholas et al., 2024) variant of a model-free Q-learning algorithm (Rescorla, 1972; Sutton & Barto, 1998). The model assumes a stored value, *Q(odor, action)*, for choosing an action of licking or not licking in response to a given odor. After each outcome, *r_t_*, the *Q* value for the chosen action was updated according to:

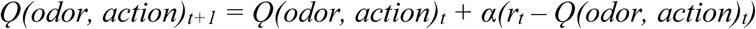

where the degree of updating was controlled by the learning rate α (**Supplementary Fig.1A**), which was a free parameter that ranges between 0 and 1. *Q* values of unchosen actions and other odors remained unchanged.

The model learned separate *Q* values for each odor (go, no-go, and control (nitrogen flow without odorant)) and action combination, such that six *Q* values were estimated in total, three for licking (*lick*) and three for not licking (*nolick*). The two *Q* values corresponding to the present stimulus, *o*, were used to compute a decision variable of the subject’s response on each trial:

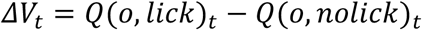

The probability of licking was modeled using a logistic function:

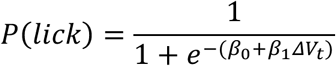

where *β*_0_ is an intercept parameter that accounts for the bias towards licking and *β*_1_ is an inverse temperature parameter that estimates the sensitivity to the learned values of actions related to the presented odor (**Supplementary Fig.1B**).

### Model Fitting

We estimated model parameters for each animal using hierarchical Bayesian inference in order to allow group-level priors to regularize subject-level estimates. This approach to fitting reinforcement learning models improves parameter identifiability and predictive accuracy (van Geen & Gerraty, 2021) and has been used to fit similar Q-learning models (Nicholas et al., 2022). We first split the subjects into two groups: one group that underwent only acquisition (*n* = 6) and another that underwent a rule reversal following acquisition (*n =* 6). This splitting was performed because we reasoned that pooling data together from all animals would artificially inflate the learning rate of animals in the acquisition-only group. This is because animals in the reversal group experienced more trials where new learning was required, effectively doubling the “volatility” of this environment (Behrens et al., 2007).

To fit the model, the joint posterior was approximated using No-U-Turn Sampling (Hoffman & Gelman, 2014) as implemented in Stan (Carpenter et al., 2017). Four chains with 2000 samples (1000 discarded as burn-in) were run for a total of 4000 posterior samples per model. Chain convergence was determined by ensuring that the Gelman-Rubin statistic, *R̂*, was close to 1.

The model’s likelihood function can be written as:

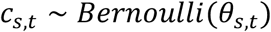

where *c_s,t_* is 1 if subject *s* chose to lick on trial *t* and 0 if the subject chose not to lick, and *θ_s,t_* is the estimated probability of this subject licking on this same trial. Following the recommendations of (Gelman & Hill, 2006), each subject’s intercept and inverse temperature *β*_s_ were drawn from a multivariate normal distribution with mean vector *μ*_*β*_ and covariance matrix *∑*_*β*_:

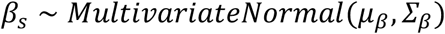

where *∑*_*β*_ was decomposed into a vector of coefficient scales *τ*_*β*_ and a correlation matrix *Ω*_*β*_ via:

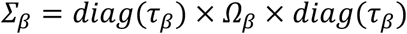

We set weakly-informative hyperpriors on the group-level hyperparameters *μ*_*β*_, *Ω*_*β*_ and *τ*_*β*_:

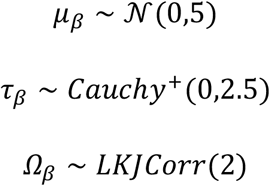

Each subject’s learning rate parameters were also fit hierarchically with the following prior and hyperpriors (*a1, a2*):

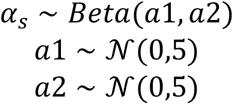

A description for why these prior and hyperpriors were chosen, as well as further details about the parameterization, can be found in Nicholas et al. (2022).

### Image acquisition and preprocessing

MRI data were acquired as previously described (Bergmann et al., 2025). In brief, scans were acquired with a 9.4 Tesla MRI (Bruker BioSpin, Ettlingen, Germany) using a quadrature 86 mm transmit-only coil (Bruker BioSpin) and a 20 mm loop receive-only coil (Bruker BioSpin), and were reconstructed using ParaVision 5.1 (Bruker). Mice underwent multiple sessions of event-related fMRI (6–14 sessions during *Acquisition*, 5–6 sessions during *Reversal*), with each session containing multiple runs (2–12 runs per session), i.e., task blocks (see Flexible discrimination learning section). The rapid-presentation event-related design was generated using Optseq2 (Greve, 2002).

Mice were anesthetized for a short period of time (5% isoflurane) at the start of each session to allow for positioning them in the scanner. For each session, a relaxation enhancement (RARE) T2-weighted structural imaging (50 coronal slices, TR/TE 2300/8.5 ms, RARE factor = 4, flip angle = 180°, 200 × 200 × 300 μm^3^, field of view of 19.2 × 19.2 mm^2^, matrix size of 96 × 96) was first acquired while the mice performed the odor discrimination task. Then, blood oxygenation-level dependent (BOLD) contrast run scans were acquired for six minutes using spin echo-echo planar imaging (SE-EPI) sequence (TR/TE 2500/13.022 ms, flip angle = 90°, 50 coronal slices, voxel size 200 ×200× 300 μm^3^, field of view of 14.4 × 9.6 mm^2^, matrix size of 72 × 48). Preprocessing of raw data included removal of the first two volumes for T1-equilibration effects, compensation for slice-dependent time shifts, rigid body correction for head motion, semi-automatic linear registration (FSL FLIRT) to the Allen Mouse Brain Common Coordinate Framework version 3 (CCFv3, Kuan et al., 2015; Lein et al., 2007) that included a manual correction step for each session to validate proper alignment, and spatial smoothing with a full width at half maximum (FWHM) of 500 μm.

### fMRI data analysis

fMRI data were analyzed using SPM12 (Wellcome Department of Cognitive Neurology, London, UK) and SnPM13 (http://nisox.org/Software/SnPM13/). The design matrices of all general linear models (GLM) computed in this study included the following nuisance regressors: global signal, ventricles signal, six motion parameters and their first-order derivatives, run constant for all runs excluding the last run, and events with frame displacement larger than the voxel size (200 μm).

### Whole-brain analysis

In order to detect brain regions involved in tracking the value of an action (lick/no-lick) in a given state, we performed a whole-brain analysis using the decision variable (see Reinforcement-learning model section) as a covariate. For each animal, all sessions of the relevant experimental stage (Acquisition/Reversal) were combined to generate a single GLM in which events were defined as stimulus onset with a duration of zero. In addition, a decision variable regressor was created as a covariant by convolving the decision variable values with the mouse hemodynamic response function (HRF) previously modeled by our group (Bergmann et al., 2025). Due to the relatively slow TR (2.5 s) and the short dynamics of the mouse HRF, it was not possible to disentangle the stimulus, choice and feedback, and therefore, events were defined as single whole trials. A linear contrast of regressor coefficients was computed at the single-subject level for the decision variable regressor and was further used for a second, group-level analysis using a one-sample *t*-test. All group-level statistical maps were corrected for multiple comparisons using family-wise error correction. For the group-level parametric analysis (*n* = 12; **Figure 2**), the voxel extension was set to 5 voxels. For non-parametric statistical maps (*n* = 6; **Figure 3**), a permutation test was performed using the sign-flip approach in which 64 permutations, the equivalent of 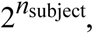 were computed. To allow for cluster-level inference, we defined the variance smoothing to be the same FWHM that was applied to the data as instructed by SnPM13 manual. The cluster-defining threshold was set to *t* statistic of 6 (∼ *p* < 0.001, df = 5, prior to cluster-level inference), resulting in a critical STCS (suprathreshold cluster size) of 3 voxels.

**Figure 2.**
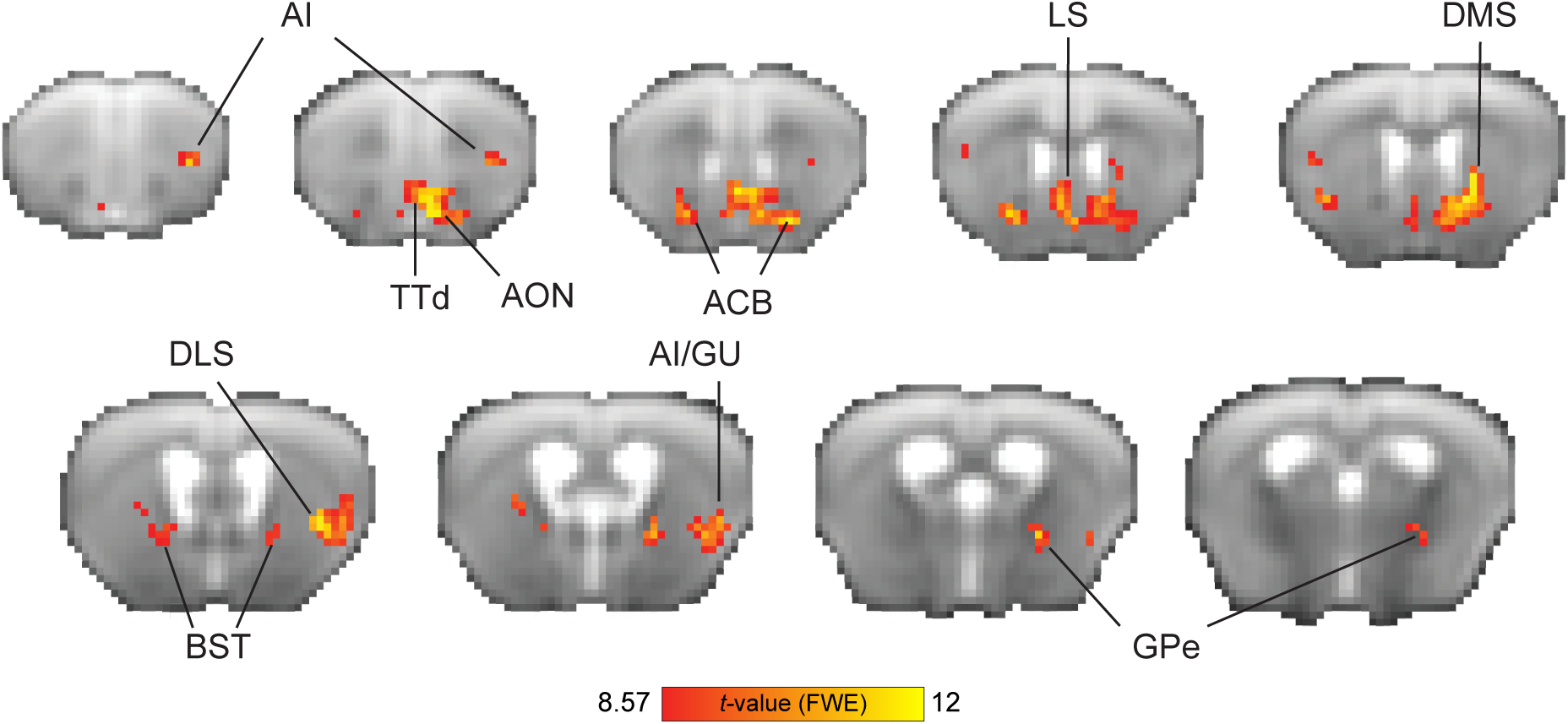
Neural substrates of Q-learning signals during go/no-go task acquisition. Group-level parametric one-sample *t* statistic maps showing BOLD response correlates of the decision variable values from the Q-learning model (*n* = 12 mice). The red color spectrum indicates areas with positive correlations to decision variable values, while blue colors indicate negative correlations. Maps are presented on the averaged raw fMRI data (spin-echo echo planar imaging) and annotated based on the Allen Mouse Brain Atlas; *p* < 0.05, corrected for multiple comparisons using family-wise error correction, voxel extension of 5. ACB, nucleus accumbens; AI, agranular insular area; AON, anterior olfactory nuclei; BST, bed nucleus of the stria terminalis; GU, gustatory cortex; DLS, dorsolateral striatum; DMS, dorsomedial striatum; GPe, globus pallidus externus; TT, tenia tecta; d, dorsal; v, ventral.

**Figure 3.**
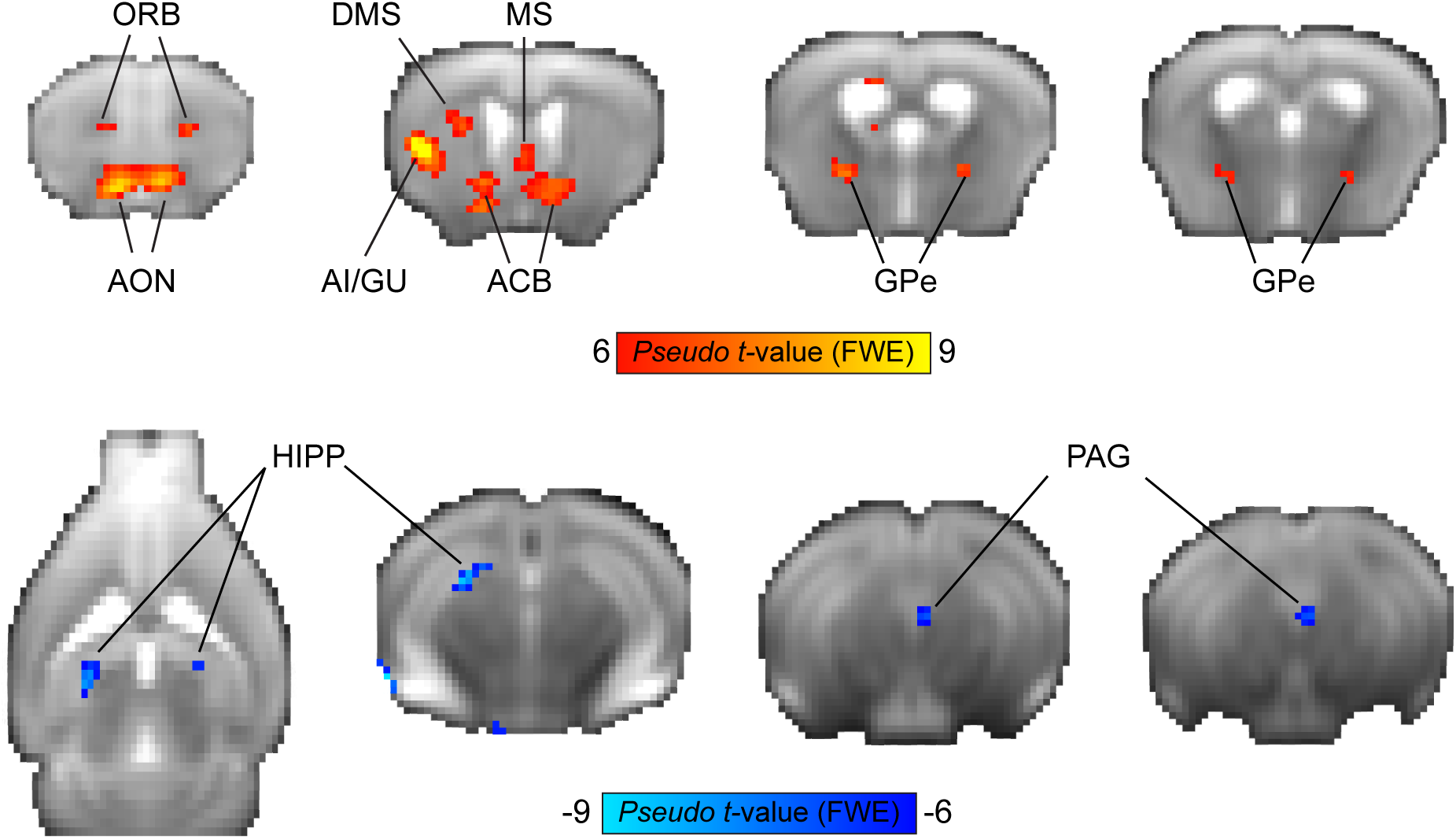
Neural substrates of Q-learning signals during reversal learning. Group-level non-parametric maps showing BOLD response correlates of the decision variable values from the Q-learning model (*n* = 6 mice). The red color spectrum indicates areas with positive correlations to the decision variable values, while blue colors indicate negative correlations. Maps are presented on an average raw fMRI data (spin-echo echo planar imaging) and annotated based on the Allen Mouse Brain Atlas; *p* < 0.05, corrected for multiple comparisons using family-wise error correction, voxel extension of 3. ACB, nucleus accumbens; AI, agranular insular area; AON, anterior olfactory nuclei; DMS, dorsomedial striatum; GPe, globus pallidus externus; GU, gustatory cortex; HIPP, hippocampus; MS, medial septum; ORB, orbitofrontal cortex; PAG, periaqueductal gray.

Brain regions identified in the whole-brain analysis but located near fiber tracts, MRI artifacts, or close to the brain’s boundaries were excluded from the results section.

### Region of interest (ROI) analysis

Following the whole-brain findings of brain regions that are correlated to the Q-learning computation, we wanted to further assess the contribution of each of these regions to specific cognitive processes. Thus, for each experimental stage (*Acquisition* and *Reversal*), we defined each trial as an event based on the subject’s response (Hit, False Alarm [FA], Correct Rejection [CR], and Miss) and generated a GLM per subject for each session separately with regressors corresponding to the different event types. We created specific ROI masks for regions identified based on the overlap between contiguous clusters of voxels in the statistical parametric/non-parametric maps and the Allen Mouse Brain Connectivity (AMBC) atlas (**Supplementary Fig. 3**). We used the MarsBaR toolbox (Brett et al., 2002) to extract finite impulse responses, plotting the hemodynamic response without assumptions on its response characteristics, with the onsets shifted by 5 s to allow to observe the pre-trial baseline and the full evolution of the response. We note that while the GLM used to identify the regions and the ROIs used in the ROI analysis are not independent, the inferences derived from this analysis are intended to characterize how Hits, FAs, CRs and Misses contribute to the response first identified in the Q-learning GLM which modeled the go and no-go conditions with the estimated decision variable as a parametric regressor. The ROI analyses therefore serve to quantify the fMRI response at both the individual-animal level and across these conditions.

### Statistical analyses

Behavioral data were analyzed using MATLAB R2023 (The Mathworks, Natick, MA, USA), except for analysis of variance (ANOVA) which was performed using jamovi 2.3. For all repeated measures ANOVA tests, time was defined as a within-subject factor and the parametric tests were corrected for sphericity violation using the Huynh-Feldt method. For fMRI data, repeated measures ANOVA was performed in R (R Core Team, 2024) using RStudio (RStudio Team, 2020), with sphericity violation corrected using the Greenhouse-Geisser method.

## Results

We utilized an experimental setup that allows for the delivery of odors into the scanner alongside a closed-loop system for lick detection and water delivery, allowing us to perform behavioral experiments with high precision in head-fixed mice (for a full description of the setup, see Bergmann et al., 2025). In this study, mice (*n* = 12) learned to perform an instrumental odor discrimination task while undergoing fMRI scanning (**Figure 1A**), with a subset of mice (*n* = 6) undergoing a rule reversal phase during which the action-outcome contingencies were switched.

### Mice learn action-outcome associations and subsequent rule reversal in a go/no-go odor discrimination task

During task *Acquisition*, mice learned to discriminate between the rewarded stimulus (S+) and the unrewarded stimulus (S-) and reached a performance criterion of >80% correct responses by the fourth day (**Figure 1B**). A repeated-measures analysis of variance (rm-ANOVA) revealed a significant effect of Time (*F*(5,11) = 25.2, *p* < 0.001, η^2^ = 0.585) as mice gradually learned the action-outcome associations for both stimuli. Evaluation of the reaction time from odor delivery to a lick response showed a decrease throughout sessions as animals became more proficient in the task (**Figure 1C**). Next, after the mice reached the criterion and maintained it, task contingencies were switched in the *Reversal* stage. Mice rapidly learned the new contingencies (**Figure 1D**), reaching a performance criterion on average within two sessions (Friedman rm-ANOVA; χ^2^ (4,6) = 16.8, *p* < 0.01). The ability to quickly adapt to the switch in contingencies can be further observed by a decrease in reaction time throughout sessions during the *Reversal* phase (**Figure 1E**). Overall, mice presented goal-directed behavior during the experiment as indicated by the gradual lick responses to rewarded stimuli only (without using punishment for incorrect responses), as can be observed by a significant difference in reaction time for the first session of reversal relative to acquisition (**Figure 1F**; Wilcoxon Signed-rank test; W **=** 20, *p* < 0.05). Importantly, when evaluating for differences between the sub-group of mice that underwent only the acquisition phase and that who underwent a following reversal phase, we found no significant differences in learning the initial action-outcome association (two-way rm-ANOVA, *F(*1,11) = 0.231, *p* = 0.947, η^2^ = 0.006).

Next, we used a reinforcement-learning model in order to parameterize the value of an action to a given odor. Specifically, we used a model-free Q-learning algorithm to allow for trial-by-trial estimation of action values (Q) of each odor to assess the process of choosing an action based on prior experiences. As estimated by the model, the probability of choosing to lick for the go odor accurately captures the experimental data (**Figure 1G**), showing a sharp decrease in probability at the point of reversal with a steep recovery. To illustrate Q-learning estimation, we plotted the decision variable values for one of the mice that participated in both *Acquisition* and *Reversal* (**Figure 1H**), demonstrating that the majority of learning occurred within approximately 500 trials in both stages (the equivalent of approximately two sessions), matching the time required to reach criterion for this animal. Similar estimation was observed for the other mice (not shown).

### Brain responses indicative of value-based decision-making in the mouse brain

We sought to characterize the neural responses that are involved in reinforcement learning during flexible discrimination learning. We used the Q-learning algorithm in order to detect brain regions that take part in computing the value of choosing an action (lick/no-lick) to a given odor stimulus. Regions that track the decision variable estimated by the model, therefore, reflect the dynamic nature of learning at the individual-animal level. Specifically, in this task, regions identified using the decision variable are implicated in learning odor-action associations: one odor signals that licking will result in water delivery, while the other odor signals that licking will not yield reward. Though incorrect responses are not explicitly punished, animals learn to avoid unrewarded actions.

To evaluate responses in the task acquisition stage, we used high-field fMRI (9.4T) to measure distributed brain activity from the naïve state to task proficiency. We entered the decision variable of each animal (*n* = 12) as a parametric modulator in first-level GLM analysis, then conducted a second-level group analysis, calculating a statistical parametric map of regions that correlate with the decision variable (**Figure 2**). We observed responses in the basal ganglia (nucleus accumbens [ACB]), dorsomedial striatum [DMS], dorsolateral striatum [DLS] and globus pallidus externus [GPe]), regions in the insular cortex (agranular insular area [AI]) related to rewards and learning the cue-reward association, regions related to odor processing (tenia tecta [TTd] and anterior olfactory nuclei [AON]), regions related to avoidance and stress regulation (bed nucleus of the stria terminalis [BST]), and regions related water consumption (posterior AI/gustatory cortex [GU]). The most prominent responses (numerically) were observed in the striatum (ACB and DMS). Overall, this measure reflects responses in brain regions previously implicated in value-based decision-making.

Next, we sought to characterize the neural responses during rule reversal. Reversal learning paradigms evaluate behavioral flexibility by switching the contingencies between stimuli and their outcomes, requiring subjects to change their learned responses when they encounter no reward for a previously rewarded response, as well as the ability to beneficially respond to a stimulus that was previously not reinforced. Using this experimental manipulation, we wanted to identify brain regions that are active during this cognitive process. Given that only a subset of mice underwent rule reversal, we used a non-parametric approach for whole-brain analysis and ran a permutation test. A GLM modeling the decision variable at the reversal stage of each animal (*n* = 6) was entered into a second-level group one-sample *t*-test, resulting in a statistical non-parametric map (**Figure 3**) of regions that were preferentially positively or negatively correlated with the value of licking or not licking, respectively. This map reveals the involvement of several regions that correlate with the decision variable values, showing a positive correlation in frontal and insular cortices (orbitofrontal [ORB] and AI/GU cortices), striatum (ACB and DMS), pallidum (medial septum [MS] and GPe) and olfactory processing regions (AON), and negative correlation in the dorsal hippocampus (HIPP) and ventral PAG.

### Activation of PAG correlates with beneficial behavioral responses in reversal learning only

Given recent findings suggesting that PAG represents an aversive prediction error, and the lack of evidence linking it to reversal learning and action value, especially under appetitive conditions, we next sought to characterize its responses in contrast to those in the ACB, a well-established region for prediction error computation. We therefore examined the PAG and ACB contributions to the different cognitive components of discrimination learning using an ROI analysis, and looked at finite impulse responses (FIR) for conditions differentiated by their behavioral responses, whether correct or incorrect, and their outcomes (Hit, Correct Rejection, False Alarm, and Miss; see **Supplementary Fig. 2** for behavioral performance).

We estimated FIR-based BOLD response time courses from the PAG and ACB and computed the mean area under the curve (AUC) across time points for each session to quantify overall response magnitude. Focusing on the PAG, a two-way repeated-measure ANOVA, with Condition (Hit, Correct Rejection, False Alarm, and Miss) and Session as within-subject factors, revealed a significant main effect of Condition (*F*(1.77, 8.87) = 15.17, *p* < 0.01, generalized η² = 0.34), indicating that AUC varied across behavioral outcomes, distinguishing correct from incorrect responses, and as can be seen in the analyses below, was driven mainly by positive responses to Correct Rejection and negative to Hit. In contrast, neither the main effect of Session nor the Session × Condition interaction was significant, suggesting that overall effects were stable across sessions, with learning occurring in the first session, consistent with the animals’ behavioral outcomes. Analysis of AUC values from the ACB showed a significant main effect of Condition (*F*(1.66, 8.30) = 20.61, *p* < .001, generalized η² = 0.55) and a significant interaction of Session × Condition (*F*(12, 60) = 2.20, *p* = .023, generalized η²= 0.19), with no significant effect of Session.

Next, we compared responses between the PAG and ACB. Given that as the animal learns the decision variable stabilizes and is maximally different between correct licks for the go odor (Hit) and avoidance for the no-go odor (Correct Rejection) (**Figure 1H**), we focused on these two conditions during the last *Reversal* session when mice had already fully learned the rule reversal (**Figure 4**). A two-way repeated-measure ANOVA revealed a significant interaction of ROI × Condition (*F* (1, 5) = 55.13, *p* < .001, generalized η² = 0.83), indicating that the behavioral effect of Condition reversed between PAG and ACB. Post hoc pairwise comparison performed within each ROI demonstrated that ACB showed a strong activation to Hit events and inactivation to Correct Rejection (**Figure 4B,C**; bottom row) (*t* (5) = 4.19, *p* = .020, Holm-adjusted). Conversely, PAG showed inactivation to Hit events and activation to Correct Rejection (**Figure 4B,C**; top row; *t* (5) = 8.95, *p* = 0.002). Additional cross-region contrasts confirmed this double dissociation, showing that for Correct Rejection the AUC was higher in PAG than in ACB, whereas for Hit the AUC was higher in ACB than in PAG (both *p* ≤ 0.025, Holm-adjusted). Taken together, these findings reveal a crossover interaction in which ACB and PAG exhibit opposite BOLD response modulations for Correct Rejection versus Hit conditions, highlighting distinct functional contributions of the two regions to outcome processing.

**Figure 4.**
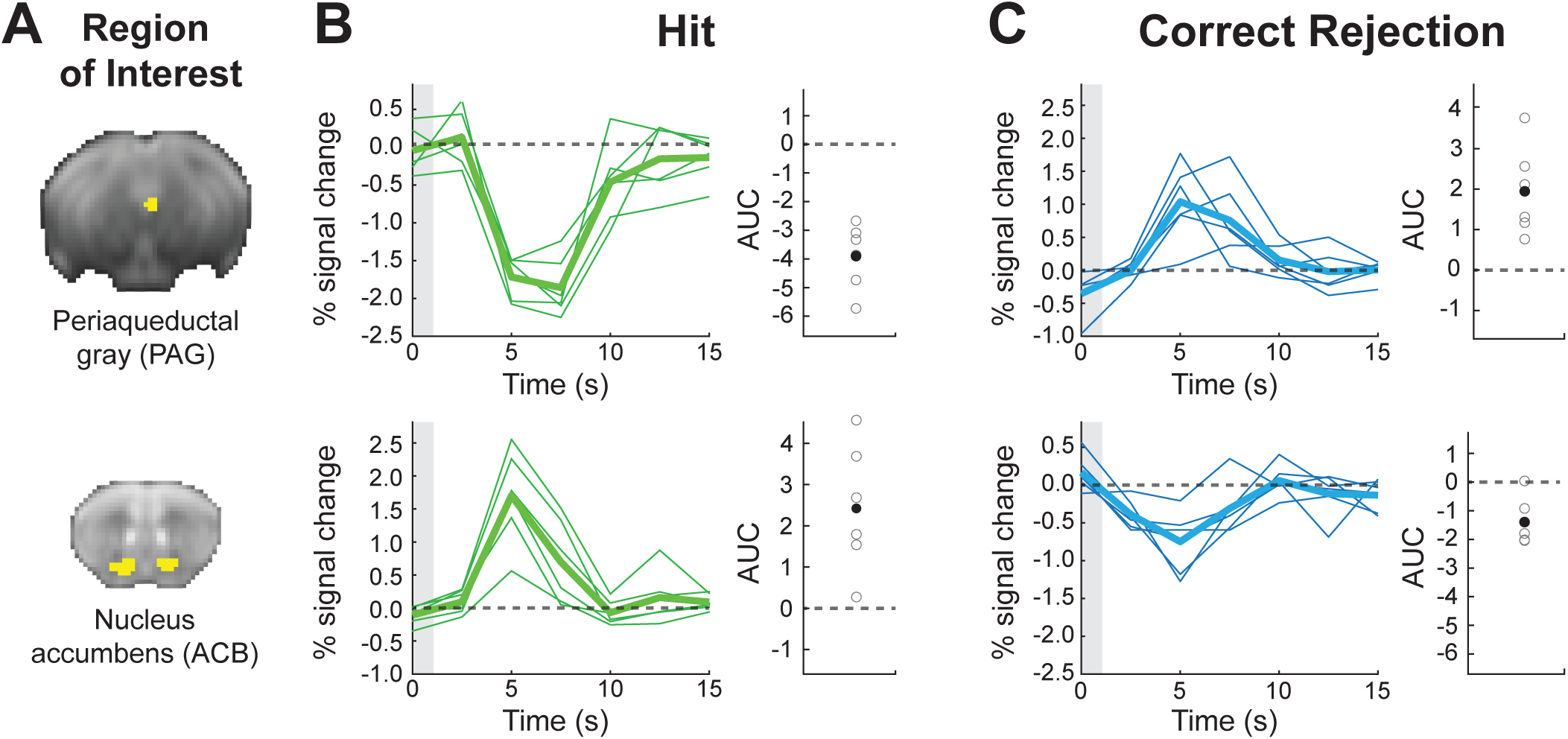
Opposing roles of the periaqueductal gray (PAG) and nucleus accumbens (ACB) in mediating adaptive behavior in proficient mice. Region of interest (ROI) analysis showing the time course fMRI BOLD response in PAG (*top*) and ACB (*bottom*) for the fifth session in the reversal stage. **(A)** ROI masks are presented on an average raw fMRI image (spin-echo echo planar imaging). **(B)** BOLD fMRI Responses to Hit (lick response to go trials; *green*) and **(C)** Correct Rejection (no lick to no-go trials; *blue*) are shown. The thick lines represent the group averaged response and the thin lines show individual animals. The gray boxes at time zero depict the odor stimulus timing (1 s). Group mean area under the curve (AUC) of the fMRI response (filled circles) and individual animals (open circles) demonstrate consistent responses at the group and individual animal levels.

Finally, we wanted to examine whether the activity observed in PAG is specific to reversal learning or whether this region contributes to acquisition learning as well (**Figure 5**). We extracted FIR responses for an *Acquisition* session at a timepoint corresponding to reversal learning, when mice demonstrated comparable performance levels (**Figure 5A**; Wilcoxon Signed-rank test; *p* = 1, Z = 0), and computed the AUC values for the two conditions. Looking at the Hit condition, we found a significant decrease in AUC values for *Acquisition* relative to *Reversal* (**Figure 5B**; Wilcoxon Signed-rank test; *p* = 0.031, Z = 21). Comparison of the Correct Rejection condition revealed a trend (**Figure 5C**; Wilcoxon Signed-rank test; *p* = 0.093, Z = 2), showing overall decrease in AUC values. Further, PAG responses during the *Acquisition* phase were not significantly different than baseline in both conditions (Sign test; Hit: *p* = 0.218, sign = 1; Correct Rejection: *p* = 1, sign = 3). Collectively, the results indicate that the subregion in PAG that was found to be important for learning of correct behavioral responses in *Reversal* does not seem to be important for *Acquisition*.

**Figure 5.**
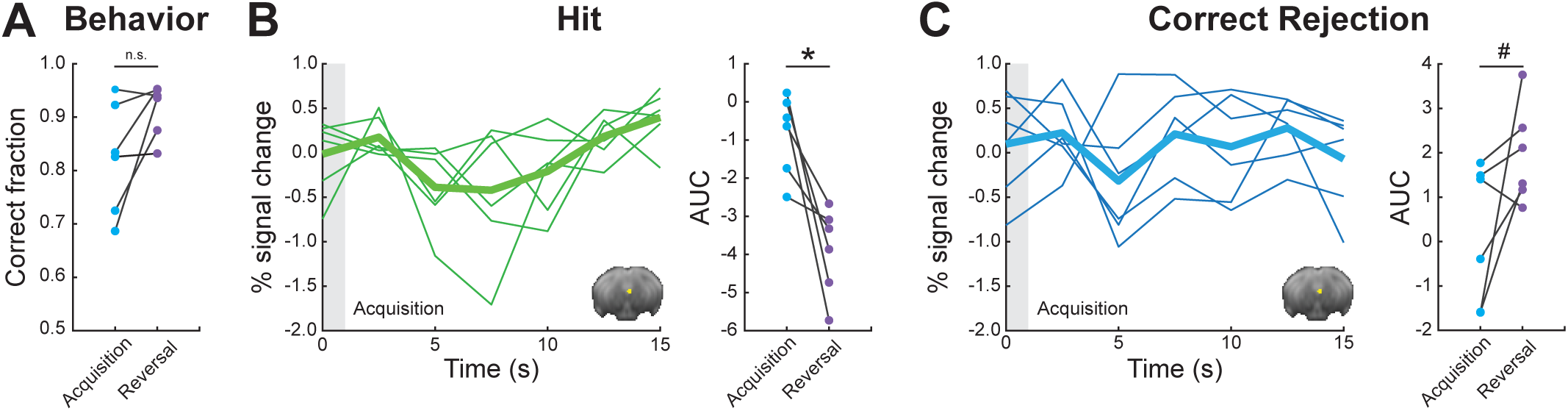
PAG responses do not contribute to adaptive behavior in proficient mice during Acquisition. **(A)** Pairwise comparison of behavioral performance during *Acquisition* vs*. Reversal* for sessions 4/5. **(B)** FIR responses for Hit condition during *Acquisition* (left). Pairwise comparison of AUC values showing individual animals for *Acquisition* vs*. Reversal* (right). **(C)** FIR responses for Correct Rejection condition during *Acquisition* (left). Pairwise comparison of AUC values showing individual animals for *Acquisition* vs*. Reversal* (right). For FIR responses, the thick lines represent group averages and the thin lines show individual animals. The gray boxes at time zero depict the odor stimulus timing (1 s). \**p* < 0.05, ^#^*p* < 0.1, n.s. no significance. Insets show PAG region of interest mask presented on an average raw fMRI image (spin-echo echo planar imaging).

## Discussion

Flexible, goal-directed behavior relies on the capacity to adapt action–outcome associations when contingencies change. This process, typically studied using reversal learning paradigms, is thought to depend on corticostriatal circuits and dopaminergic teaching signals, yet it remains unclear whether additional brain regions may also contribute. Whole-brain imaging approaches provide an opportunity to address this question. In this study, we found that the periaqueductal gray (PAG) contributes to cognitive flexibility by supporting the suppression of previously reinforced actions in the absence of explicit punishment, and assessed its contributions relative to the nucleus accumbens. By combining task-based fMRI with reinforcement learning modeling in mice performing an odor discrimination task, we found that acquisition was supported by the nucleus accumbens, whereas reversal learning additionally engaged the PAG. Notably, PAG responses contrasted with those in the nucleus accumbens, exhibiting preferential activation during correct rejection of no-go cues and suppression during correct approach to go cues. Together, these results identify the PAG as a key contributor to reversal learning, expanding current models of the neural mechanisms underlying behavioral flexibility. Moreover, they highlight the strength of whole-brain fMRI in rodents, demonstrating that novel findings can emerge even within a well-established and extensively studied behavioral paradigm.

By applying a model-free reinforcement-learning algorithm we were able to capture trial-by-trial dynamics of value updating and link them to fMRI responses. This computational approach was essential for detecting the emergence of PAG activity during reversal, as it allowed us to model how action values evolve across individual trials rather than relying on averaged performance measures. Computational modeling of reinforcement learning has been highly influential in human fMRI studies (Niv, 2009; O’Doherty et al., 2003), and its value has only recently been shown in rodent fMRI as well (Winkelmeier et al., 2022). Our results demonstrate the feasibility and utility of such models in animal neuroimaging, highlighting how model-based analyses can improve sensitivity to dynamic neural processes underlying learning. This methodological advance also paves the way for more direct cross-species comparisons of reinforcement learning circuitry.

Beyond the striatum, our whole-brain analyses revealed that acquisition engaged a distributed set of regions, including the agranular insula, gustatory cortex, anterior olfactory nuclei, and bed nucleus of the stria terminalis. These findings are consistent with prior reports that value-based learning recruits not only canonical reward regions but also areas involved in sensory processing, interoception, and stress regulation (FitzGerald et al., 2013; Ge & Balleine, 2022; Hernández-Ortiz et al., 2023; Kogan & Fontanini, 2024; Levinson et al., 2020). The engagement of olfactory and gustatory cortices likely reflects the multimodal nature of the task, while activity in the insula and bed nucleus of the stria terminalis may reflect the integration of reward signals with internal state and arousal. Together, these observations emphasize that flexible discrimination learning emerges from the interaction of distributed brain systems, with the PAG contributing selectively during reversal to bias action suppression once initial associations have been formed.

Classically, the PAG has been implicated in defensive behaviors (Bandler & Keay, 1996; Carrive, 1993), processing nociceptive signals (Basbaum & Fields, 1984; Behbehani, 1995), and coordinating autonomic responses to threat (Dampney, 1994; Keay & Bandler, 2001). However, there has been emerging evidence linking it to behavioral flexibility and value-based decision-making (Ozawa et al., 2017; Reis et al., 2021; Sukikara et al., 2006; Wright & McDannald, 2019). Recent work has highlighted PAG as a potential relay between brainstem value signals and forebrain decision circuits (Gorka et al., 2023; Roy et al., 2014). Further, PAG neurons were shown to encode both negative and positive prediction errors (Walker et al., 2020; Wright & McDannald, 2019), and project to thalamic and cortical areas implicated in strategy updating (Assareh et al., 2016; Faull et al., 2019; Kragel et al., 2019; Krout & Loewy, 2000). Thus, PAG serves as a hub that transforms aversive sensory input into adaptive motor and physiological outputs, thereby guiding rapid survival-related responses. While the majority of previous studies linking PAG to reinforcement learning processes used aversive, pain-related paradigms, our task design allowed for reversal learning to occur in the absence of explicit punishment. Mice adapted their behavior solely through the omission of expected reward, suggesting that PAG activity may contribute to updating value representations when contingencies change, even under neutral conditions. Our results extend the current framework by showing that PAG recruitment can occur in appetitive tasks without negative reinforcement, highlighting its broader role in signaling the need to suppress outdated responses. This observation aligns with a proposal that PAG contributes to the evaluation of approach versus avoidance strategies in changing environments (Tryon & Mizumori, 2018) and suggests it may act as a general mediator of adaptive response suppression.

The region of interest analysis revealed a striking double dissociation between PAG and nucleus accumbens activity during reversal learning. Whereas nucleus accumbens responses were strongest for rewarded licks (hits), PAG responses were selectively enhanced during correct rejections. This opposing response indicates that PAG and nucleus accumbens contribute complementary signals to support flexible decision-making. The nucleus accumbens has long been implicated in representing positive prediction errors and guiding approach behavior (Montague et al., 1996; Nicola, 2007; Schultz et al., 1997). By contrast, the PAG response profile suggests a role in reinforcing the suppression of actions that no longer yield reward, consistent with its established involvement in aversive learning (Johansen et al., 2010; Walker et al., 2020). This opponency suggests that midbrain and striatal circuits jointly encode both the drive to exploit rewarded contingencies and the need to avoid perseverating on outdated responses. Through its reciprocal connections with both brainstem neuromodulatory centers and midbrain dopamine neurons, the PAG is anatomically poised to influence the teaching signals that drive reinforcement learning even in conditions beyond those classically associated with it. Namely, avoidance learning could, at least in part, be mediated by PAG computations by biasing dopaminergic signaling toward actions in situations where no explicit punishment occurs.

Importantly, PAG responses were not observed during initial acquisition, even when behavioral performance was comparable to that achieved in reversal. This indicates that PAG recruitment is not a general feature of value-based learning, but rather emerges selectively when animals must overcome prior learning. The specificity of PAG involvement in reversal echoes prior rodent and primate studies implicating cortical circuits, the orbitofrontal cortex in particular, in behavioral flexibility (Cools et al., 2002; Ghahremani et al., 2010; Izquierdo et al., 2017; Schoenbaum et al., 2006). Our findings extend this literature by demonstrating that the PAG is also selectively engaged under reversal conditions. This supports the notion that flexible decision-making depends on coordinated contributions from both cortical and subcortical regions, with PAG providing a key midbrain computation to facilitate behavioral adaptation.

Collectively, our results expand the functional repertoire of the PAG beyond its established role in aversion and defensive behaviors, positioning it as a key node in the neural circuitry that supports flexible decision-making. By demonstrating that PAG activity is selectively recruited during reversal learning, and exhibits functional opponency with striatal reward signals, our findings suggest that PAG contributes to updating value representations when prior contingencies are no longer valid. This role is particularly notable given that mice adapted their behavior without explicit punishment, implying that PAG computations may bias action selection toward adaptive avoidance even in neutral contexts. More broadly, these results underscore the importance of brainstem–forebrain interactions in reinforcement learning and provide a systems-level framework for future studies examining how PAG signals integrate with dopaminergic and cortical circuits to support cognitive flexibility. Elucidating these mechanisms will be critical for understanding how distributed midbrain circuits contribute to adaptive behavior and how their dysfunction may contribute to neuropsychiatric disorders characterized by impaired behavioral flexibility.

## Acknowledgments

This work was supported by the Israel Science Foundation (770/17; to I.K.), the National Institutes of Health (1R01NS091037; to I.K.), the Adelis Foundation (to I.K.) and the Prince Center (to I.K.). We thank Technion’s Biological Core Facilities and Edith Suss-Toby for her assistance with the MRI, the Technion Preclinical Research Authority and Nadav Cohen for assistance with animal care, Yael Niv for her helpful comments on the manuscript, and Matteo Farinella for his assistance in illustrating the behavioral setup shown in Figure 1A. We acknowledge the use of generative AI tools (ChatGPT, GPT-4.5, Open AI) to assist in manuscript editing, specifically for grammar and style improvements, as well as enhancing overall readability. These tools did not influence the scientific content or interpretation of our data. Conflict of Interest: None declared.

## Author contributions

D.L. and I.K. designed the study. D.L. and E.B. conducted experiments. D.L., E.B. and J.N. wrote software for data analysis. R.T.G. and D.L. designed fMRI data analysis. D.L. analyzed the data. D.L. and I.K. prepared the manuscript.

## Data availability

Raw imaging data and all statistical maps included in this report will be made available in BIDS format on OpenNeuro upon publication.

## Supplementary Materials

**Supplementary Figure 1.**
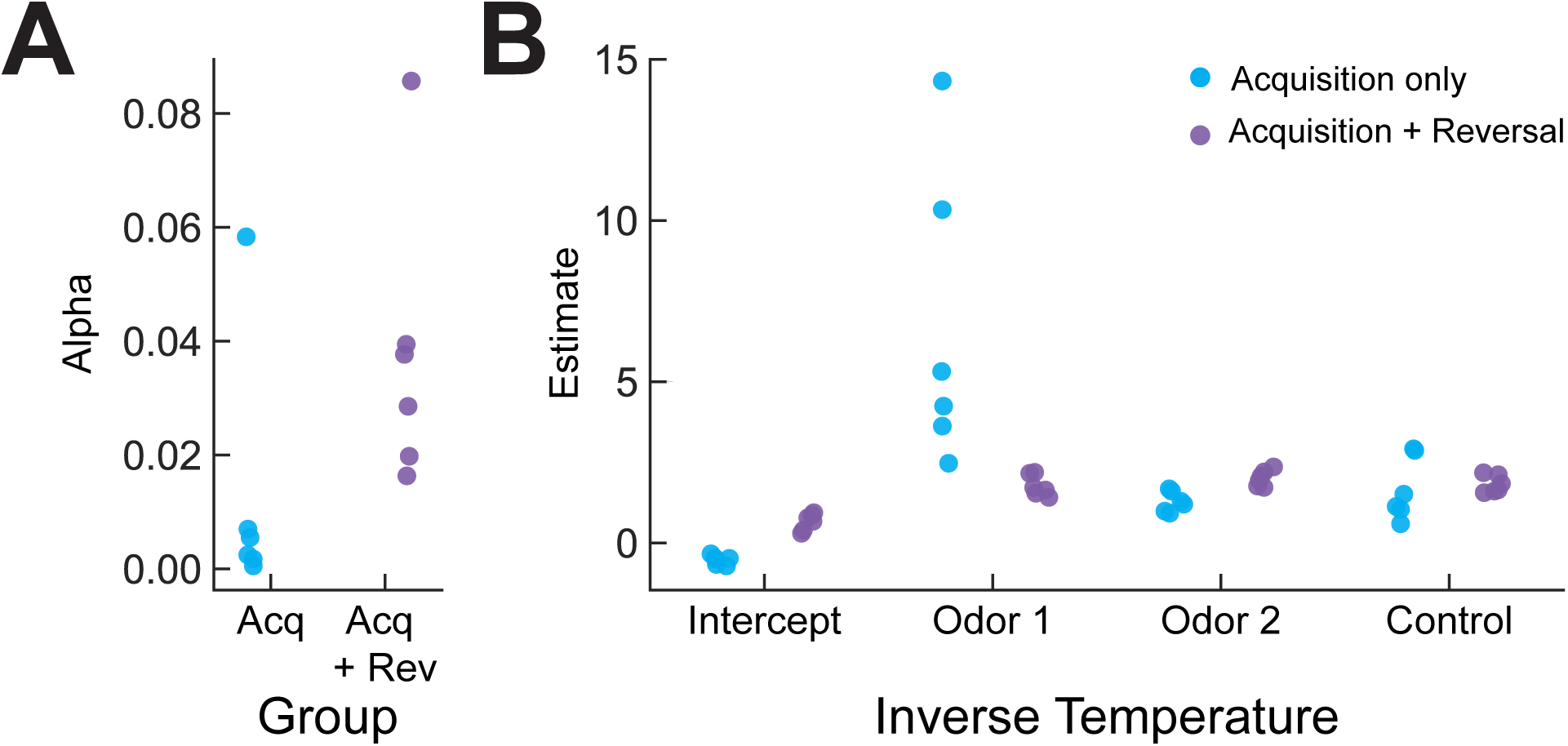
Group parameters computed by the Q-learning algorithm. Fitting parameters are shown for the two modeling groups: acquisition only (*blue*) and acquisition following reversal (*purple*; *n* = 6 per group). **(A)** Learning rate parameter alpha. **(B)** Beta estimates for intercept (bias to lick), odor 1 (go stimulus), odor 2 (no-go stimulus) and control for changes in airflow during final valve opening (nitrogen).

**Supplementary Figure 2.**
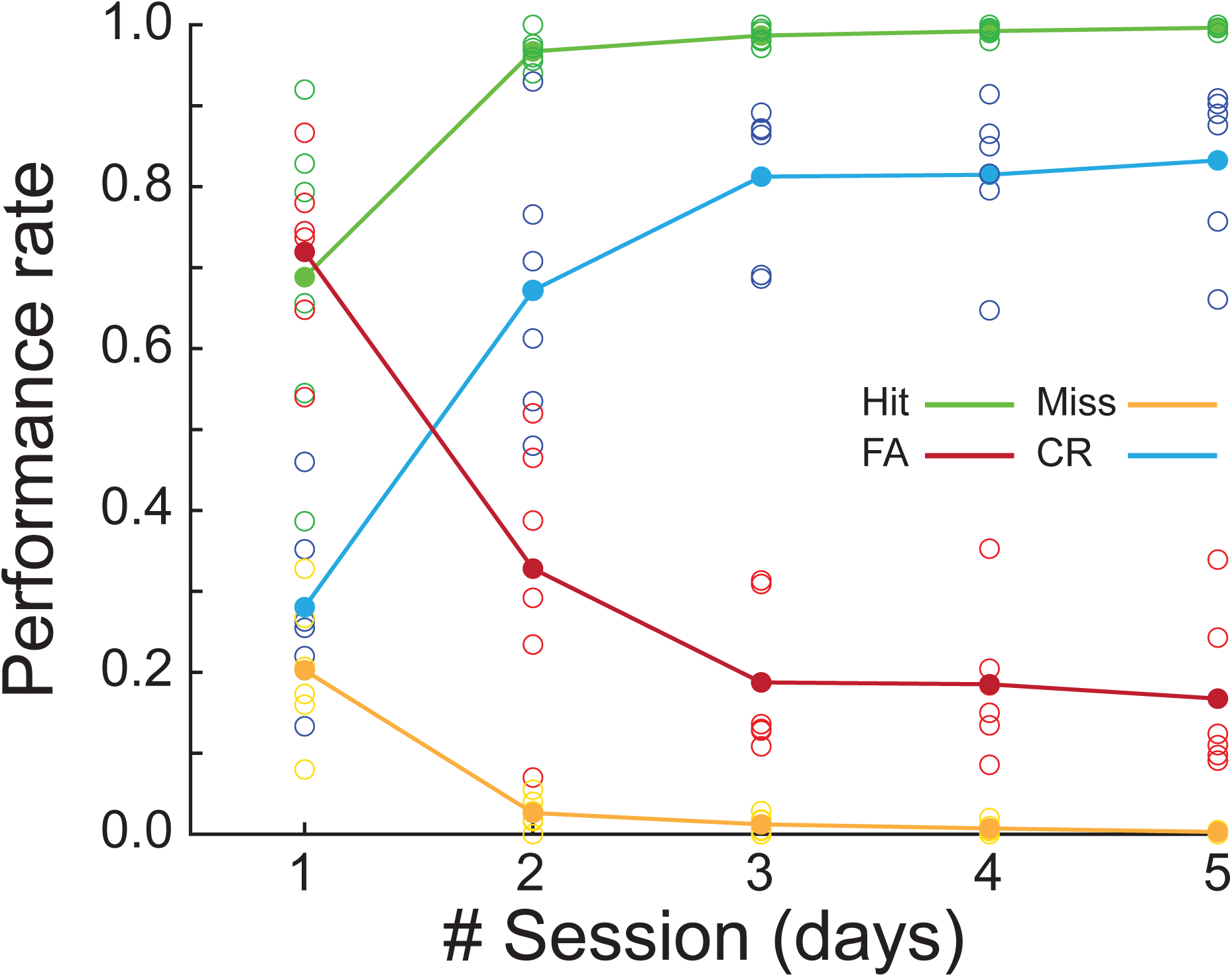
Group performance for the different discrimination conditions during Reversal. Behavioral performance rates across five consecutive sessions during the Reversal phase divided by the lick response of the animal to each cue type: Hit (lick to go odor; *green*), Miss (no lick to go odor; *yellow*), False Alarm (FA, lick to no-go odor; *red*) and Correct Rejection (CR, no lick to no-go odor; *light blue*). Filled circles indicate group mean and open circles indicate individual subjects (*n* = 6).

**Supplementary Figure 3.**
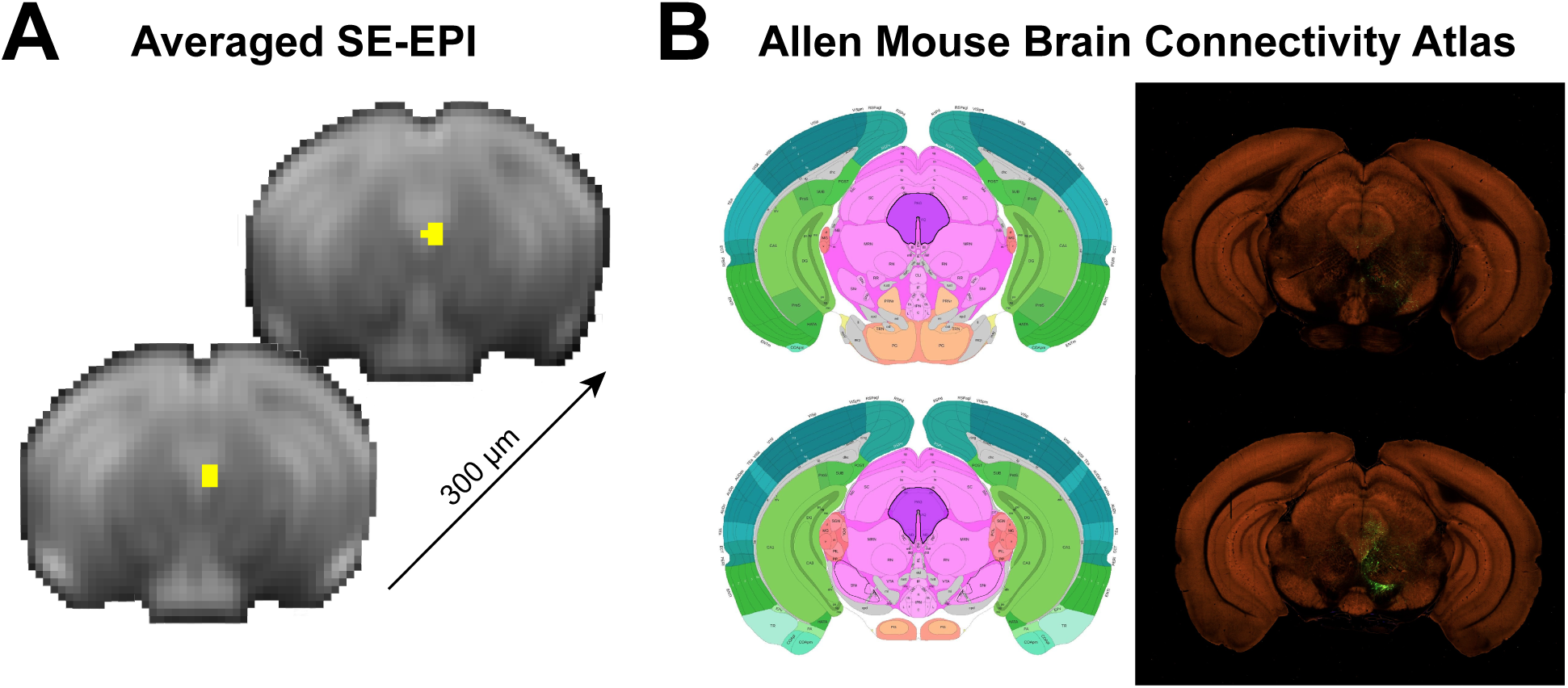
Spatial localization of the periaqueductal gray across fMRI and atlas space. **(A)** Periaqueductal gray mask (yellow) defined based on regions showing a significant BOLD response in the whole-brain analysis and further used in the ROI analysis. The ROI mask is presented on an average raw fMRI data (spin-echo echo planar imaging), shown as sequential coronal slices with a slice thickness of 300 µm. **(B)** Coronal images (*left*, atlas; *right*, two-photon tomography) taken from the Allen mouse brain connectivity atlas that correspond to the spatial location of the fMRI data. The region highlighted in purple denotes the Periaqueductal gray as defined by the Allen Institute. Image identification numbers are 87 (bottom) and 90 (top) in the reference atlas.

**Supplementary Table 1.** Summary of number of sessions completed by each subject.

| Subject | # Sessions |  |  |  |
| --- | --- | --- | --- | --- |
|  | Behavior |  | fMRI |  |
|  | *Acquisition* | *Reversal* | *Acquisition* | *Reversal* |
| *Subject 01* | 10 | — | 9 | — |
| *Subject 02* | 10 | — | 9 | — |
| *Subject 03* | 6 | — | 6 | — |
| *Subject 04* | 9 | 5 | 8 | 5 |
| *Subject 05* | 15 | — | 15 | — |
| *Subject 06* | 16 | 6 | 14 | 6 |
| *Subject 07* | 15 | 6 | 13 | 6 |
| *Subject 08* | 14 | 6 | 12 | 6 |
| *Subject 09* | 9 | — | 9 | — |
| *Subject 10* | 9 | — | 9 | — |
| *Subject 11* | 8 | 5 | 8 | 5 |
| *Subject 12* | 8 | 5 | 8 | 5 |

Summary of maximal number of sessions (one session per day) completed by each subject during the *Acquisition* and *Reversal* experimental phases (Behavior). All subjects reached learning criterion in the *Acquisition* phase by the fourth session ( $3.583 \pm 1.621$ , mean ± SD). Of the 6 subjects participating in the *Reversal* phase, 3 subjects completed 8–9 sessions, and 3 completed 14–16 sessions. This manipulation was used to rule out that an extended number of sessions during *Acquisition* affects the results observed during *Reversal*. The table also shows the number of usable fMRI data (fMRI) session, as some were excluded due to software issues or poor performance inside the scanner.

## Notes

### Competing Interest Statement

The authors have declared no competing interest.

### Summary of Updates

The abstract in the originally uploaded PDF contained an outdated version of the text, differing from the correct version in three places: *Sentence 3* PubMed: "Widely used in human research, whole-brain imaging in rodents has been a major challenge that has only recently been properly addressed." PDF: "Widely used in human research, the challenge of functional fMRI in rodents has only recently been properly addressed, opening the door to whole-brain imaging of behaving mice." *Sentence 4* PubMed: "Using functional MRI in behaving mice performing a go/no-go odor discrimination task, we compared..." PDF: "Here, we used functional MRI in mice performing a go/no-go odor discrimination task, we compared..." *Sentence 5* PubMed includes an explicit framing sentence before the trial-by-trial estimates: "To link neural activity to underlying learning processes, we modeled value updating using a model-free reinforcement-learning algorithm." Preprint omits this sentence entirely, jumping directly to: "Trial-by-trial estimates of state-action values from a model-free reinforcement-learning algorithm allowed us to dissociate..."

